# Alcohol preferring P rats exhibit aversion resistant drinking of alcohol adulterated with quinine

**DOI:** 10.1101/689919

**Authors:** Nicholas M. Timme, David Linsenbardt, Maureen Timm, Taylor Galbari, Ethan Cornwell, Christopher Lapish

**Affiliations:** Indiana University – Purdue University Indianapolis; University of Utah

## Abstract

Understanding why some people continue to drink alcohol despite negative consequences and others do not is a central problem in the study of alcohol use disorder (AUD). In this study, we used alcohol preferring P rats (a strain bred to prefer to drink alcohol, a model for genetic risk for AUD) and Wistars (control) to examine drinking despite negative consequences in the form of an aversive bitter taste stimuli produced by quinine. Animals were trained to consume 10% ethanol in a simple Pavlovian conditioning task that paired alcohol access with an auditory stimulus. When the alcohol was adulterated with quinine (0.1 g/L), P rats continued to consume alcohol+quinine at the same rate as unadulterated alcohol, despite a demonstrated aversion to quinine adulterated alcohol when given a choice between adulterated and unadulterated alcohol in the home cage. Conversely, Wistars decreased consumption of quinine adulterated alcohol in the task, but continued to try the alcohol+quinine solution at similar rates to unadulterated alcohol. These results indicate that following about 8 weeks of alcohol consumption P rats exhibit aversion resistant drinking. This model could be used in future work to explore how biological basis of alcohol consumption and genetic risk for excessive drinking lead to drinking that is resistant to devaluation.

## Introduction

Drinking in spite of the negative consequences characterizes advanced stages of an alcohol use disorder (AUD) (Sanchis-Segura & Spanagel, 2006). At this stage of an AUD, drinking can become inflexible to the point where the individual is unable to abstain in spite of negative social, legal, and health consequences. This stage of drinking is typically referred to as compulsive or aversion-resistant drinking (Hopf, Chang, Sparta, Bowers, & Bonci, 2010; Hopf & Lesscher, 2014) and is of particular concern because treatments that incorporate aversive stimuli (e.g., disulfiram) may be substantially less effective at this stage of the disorder. Genetic and environmental factors strongly influence the progression of this disorder from social to problem drinking (Edenberg & Foroud, 2013; Enoch, 2013; Field & Cox, 2008; Kreusch, Vilenne, & Quertemont, 2013; Wiers et al., 2014). The goal of the current study was to assess the impact of genetic risk on a specific form of aversion resistant drinking (quinine resistant drinking) in a rodent model of genetic risk (alcohol preferring P rats) and a control rodent strain (Wistars) during a simple task with alcohol-paired stimuli.

It is believe that a large component of risk for an AUD is genetic and human work has clearly outlined several genetic factors that are associated with the risk for an AUD (Edenberg & Foroud, 2013; Enoch, 2013). Selective breeding procedures have provided an effective way to assess the heritable aspects of AUDs and provide clear support for the role of genetics in the transmission of excessive drinking phenotype from parents to progeny. Several rodent lines are available that have been selected on different features of alcohol preference. The alcohol preferring (P) rat is line that has been selected for home cage ethanol drinking (Bell, Rodd, Lumeng, Murphy, & McBride, 2006; McBride, Rodd, Bell, Lumeng, & Li, 2014). This line is used extensively as a rodent model of AUD as they meet several criteria for a rodent model of AUD. Furthermore, they also exhibit alterations in cognitive behaviors that model some aspects of the human condition (Beckwith & Czachowski, 2014, 2016; Linsenbardt, Smoker, Janetsian-Fritz, & Lapish, 2016). These changes in behavior are likely mediated by numerous biological and physiological abnormalities observed in these animals (Engleman, Ingraham, McBride, Lumeng, & Murphy, 2006; Gilpin, Stewart, & Badia-Elder, 2008; Zhou et al., 2013). In the current study, we have chosen to use the P rat due to its high drinking phenotype and genetic load for excessive drinking.

In addition to genetic risk, experiences with and environment exposure to alcohol-paired stimuli have also been shown to play a key role in AUD. Stimuli associated with alcohol acquire incentive motivational properties that are capable of inducing craving and alcohol seeking behaviors (Field & Cox, 2008; Kreusch et al., 2013; Wiers et al., 2014). Therefore, in this study we chose to use a Pavlovian conditioning task to administer alcohol following alcohol-paired auditory stimuli (Linsenbardt & Lapish, 2015; Linsenbardt, Timme, & Lapish, 2018; McCane, Czachowski, & Lapish, 2014). By training animals to consume alcohol in this task, we were able to examine task acquisition and performance in a model of genetic risk (P rats) and controls (Wistars).

After training in this task, quinine (a bitter tasting, aversive substance) was added to the alcohol and drinking was assessed across both populations of animals. Continued drinking despite the presence of quinine has been used previously as an assessment of aversion-resistant drinking (Hopf et al., 2010). Wistars (control) showed a substantial decrease in consumption upon quinine adulteration during the task, while P rats (model for genetic risk) did not. This was despite that fact that the same P rats preferred non-quinine adulterate alcohol over quinine adulterated alcohol in their home cage. Furthermore, Wistars continued to try the quinine adulterated alcohol throughout the test session, but their overall consumption significantly decreased. These results indicate that the Wistar rats maintained flexible control over drinking, despite their continued motivation to drink, whereas P rats did not.

## Methods

### Animals

This study utilized 12 male alcohol preferring (P) rats supplied by Indiana University School of Medicine and 12 male Wistar rats (Envigo, Indianapolis). All animals were born between November 22-26, 2017. Both sets of animals were shipped via ground transportation from breeding facilities to the laboratory, all within the city of Indianapolis. All animal procedures were approved by the Indiana University Animal Care and Use Committee.

### Intermittent Access Protocol

An intermittent access protocol (IAP) was used to acclimate animals to the taste and effects of alcohol (Linsenbardt & Lapish, 2015; Simms et al., 2008). Free access to 20% ethanol in one bottle and tap water in another bottle was provided in the animals’ home cages for 24 hour periods on alternating days. Animals were weighed and bottles were placed on the cages on Monday, Wednesday, and Friday mornings (approximately 2 hours into the animals’ dark cycle) for 4 weeks. Bottles were pulled 24 hours later and weighed to assess consumption.

### Auditory 2CAP Task

All training and testing was conducted in standard rat shuttle boxes (Med Associates) which were housed in custom sound attenuating chambers. One retractable sipper and one speaker were located on both ends of the chamber. Only 10% ethanol solution was used throughout all training and testing in the 2CAP. An open guillotine door separated the two sides of the chamber. The chamber was illuminated by house lights located behind the chamber. Licks were recorded using electrical contacts between the sipper and the metal grid floor. Infrared photo beams were used to record the animal’s position in the chamber. Testing software was implemented in Med Associates (see Supplemental for software). A similar visual 2CAP task has been described previously (Linsenbardt & Lapish, 2015; Linsenbardt et al., 2018; McCane et al., 2014).

During the first three days of regular 2CAP task training, a short task was conducted to acclimate the animals to the operant chambers and motorized sippers prior to the 2CAP task. At the start of the acclimation task, the house lights would turn on and both sippers would be inserted into the chamber. After 20 seconds, the sippers would withdraw for 2 seconds and then be reinserted into the chamber. If, at any point, the animal reached 15 licks on a sipper, that sipper would withdraw and not re-enter the chamber. Once the lick limits had been reached on both sippers, the house lights would turn off and the acclimation session would be complete. If the lick limit on both sippers was not reached in 6 minutes, the lights would turn off, the sippers would withdraw, and the acclimation session would be complete.

All 2CAP training and testing sessions consisted of 60 trials with the house lights on. Each trial was one of three types: stay, go, or null. Stay trials were those in which the sipper would be inserted on the same side of the chamber as the animal. Go trials were those in which the sipper would be inserted on the opposite side of the chamber as the animal. Null trials were those in which the sipper would not be inserted.

The sequence of events in a trial are shown in Figure 1 A. A trial began with a 2 second attention tone of 8 kHz. Following the attention tone, a trial type tone (4, 8, or 12 kHz) was played to direct the animal to the correct access location. All null trials were associated with the 8 kHZ tone, while the 4 and 12 kHz tones were associated with stay and go trials in a counterbalanced fashion. Following the direction tone, both sippers were inserted into the chamber half-way. The correct sipper continued into the chamber, while the incorrect sipper (or both sippers in the case of a null trial) was withdrawn. Both sippers were initially inserted to prevent animals from relying on sipper sounds to locate access.

**Figure 1:**
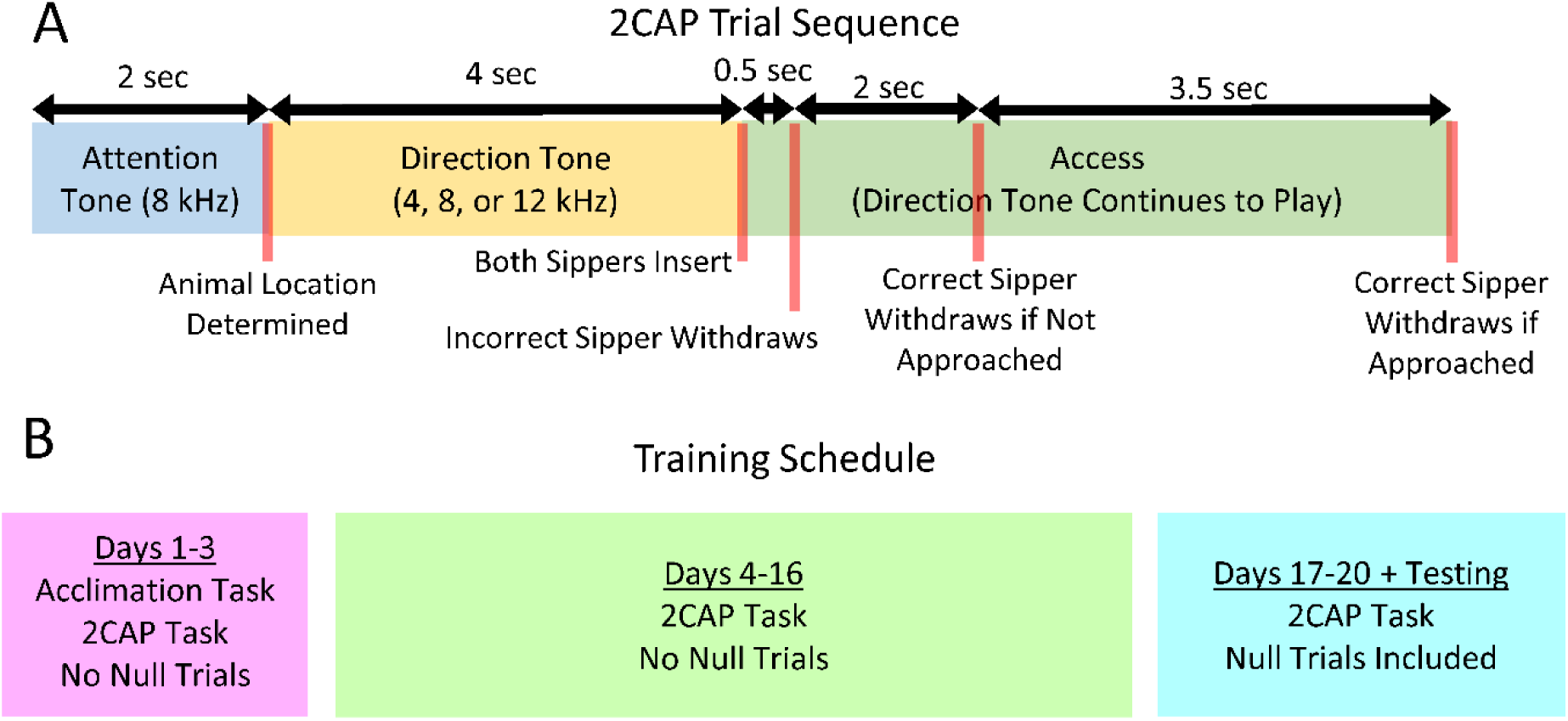
2CAP trial sequence and training schedule. **(A)** 2CAP trial sequence. A 8 kHz attention tone was played for 2 seconds, followed by a 4 second direction tone. Next, both sippers were inserted and 0.5 seconds later (half the total time of complete sipper insertion), the incorrect sipper was withdrawn (or both sippers were withdrawn in null trials). If the animal did not approach the correct sipper after 2 more seconds, it was withdrawn to prevent correcting. If the animal approached the correct sipper, it was withdrawn after a total of 5.5 seconds of access. **(B)** Training schedule. Days 1-3: The acclimation task was performed immediately before the 2CAP task with no null trials. Days 4-16: The 2CAP task was performed with no null trials. Days 17-20 and Testing (quinine and reversal): The 2CAP task was performed with null trials.

Following sipper insertion, animals were given 5.5 seconds of access to the alcohol solution. However, if the animal did not approach the sipper (i.e., break the photo beam in front of the sipper), the sipper was withdrawn after 2 seconds of access to prevent moving over to the opposite sipper (e.g. correcting). After the access period, the direction tone was turned off, the correct sipper withdrew (in the case of non-null trials where it was approached), and an inter-trial interval with no stimuli was applied prior to the next trial. The inter-trial interval period was chosen pseudorandomly from 23, 29, 31, 37, 41, 43, 47, and 53 seconds. Also, a 3 second buffer delay was used at the start and end of each trial.

To prevent animals from perseverating on stay or go trials, force trials were imposed such that if an animal drank on only one type of trial for three trials in a row, subsequent trials of the opposite type would be imposed until the animal drank on the other type of trial. For instance, if an animal drank on three stay trials in a row, it would receive go trials in place of any subsequent stay trials until it drank.

Training sessions were conducted once a day during the animals’ dark cycle on week days (Figure 1 B). On training days 1-3, the animals performed the acclimation task immediately before the 2CAP task with no null trials (30 stay and 30 go trials). For days 4-16 of 2CAP training, the acclimation task was not performed and the 2CAP task contained no null trials. From the 17^th^ day of 2CAP training onwards (including quinine and reversal testing), the animals received 20 stay, 20 go, and 20 null trials.

### Quinine Testing

The 10% alcohol solution was adulterated with 0.1 g/L quinine in a regular 2CAP session to assess aversion resistant drinking. This testing session immediately followed (1 day later) a regular 2CAP session with no quinine, which served as a baseline measurement of the animal’s behavior in the 2CAP task.

### 3-Bottle Choice Testing

A 3-bottle choice test was conducted in the animals’ home cages to ensure that P rats found the 0.1 g/L dose of quinine aversive. Each animal was given 3 days of continuous access to 2 bottles with 10% ethanol and 1 bottle with tap water. Consumption was measured once a day by weighing bottles. Following the 3 days of baseline testing, the alcohol bottle that was preferred by the animals over the course of all 3 days was adulterated with 0.1 g/L quinine. The preference for the quinine adulterated bottle was then measured after 24 hours of access. In both cases, preference was simply calculated as the ratio of alcohol consumed in 1 bottle to the total alcohol consumed from both bottles.

### Reversal Testing

To assess the degree to which animals were using the directional tones to locate ethanol access, a tone reversal test was conducted. During this test, the relationship in tone frequency between stay and go tones was reversed. For instance, animals that had learned to associate 4 kHz with go and 12 kHz with stay were instead presented with 12 kHz as the go signal and 4 kHz as the stay signal. All other features of the task were maintained. This testing session immediately followed (1 day later) a regular 2CAP session with the standard trial type/tone frequency relationship that the animal had learned. This previous day served as a baseline measurement of the animal’s behavior in the 2CAP task.

### Free Access

Near the end of testing, animals were given a free access session in which the sippers were inserted throughout the entire time of the 2CAP session. No tones were played and the house lights were on. This test was conducted to assess the animals’ motivation to drink 10% ethanol during the regular 2CAP session.

### Blood Ethanol Concentration (BEC) Measurements

Blood samples were taken from the tip of the tail to measure blood ethanol concentration (BEC) immediately after the free access session and another regular 2CAP session. Blood samples were centrifuged, blood plasma was collected and stored at −80 until analysis. Blood plasma was thawed and then run through an alcohol analyzer (Analox) to determine BEC of the blood sample.

### Training and Testing Schedule

All animals were 85 to 89 days old at the start of IAP. Animals were divided into two cohorts following IAP (4 Wistars and 4 P rats in each cohort). The first cohort entered training immediately after IAP (age: 122 days). The second cohort entered training after a delay of several weeks (age: 155 to 159 days). Testing occurred in a set order of quinine, reversal, 3-bottle choice, free access, and regular 2CAP BEC check. Some animals proceeded to testing immediately following training. Other animals were delayed to simulate the schedule following surgery.

### Statistics, Data, and Software

Statistical testing (ANVOA) and linear regression fits were performed in MATLAB and are shown in the supplemental code. All data is freely available as supplemental material, along with MATLAB code used in the analysis and the Med Associates software used to run the 2CAP task and tests.

## Results

### Intermittent Access Protocol

An intermittent access protocol (IAP) was used to acclimate animals to the taste of alcohol (Figure 2 A and B). This procedure involved 24-hour access periods to 20% ethanol and water on alternating days (M, W, F) for 4 weeks. Alcohol intake increased throughout IAP (main effect of IAP session, F(11,242) = 15.821, p < 10^−6^). Also, P rats consumed more alcohol than Wistars (main effect of strain, F(1,22) = 78.537, p < 10^−6^). Finally, the interaction between strain and IAP session was found to be significant (F(11,242) = 5.18, p < 10^−6^), indicating that the escalation in drinking through IAP was different between P rats and Wistars.

**Figure 2:**
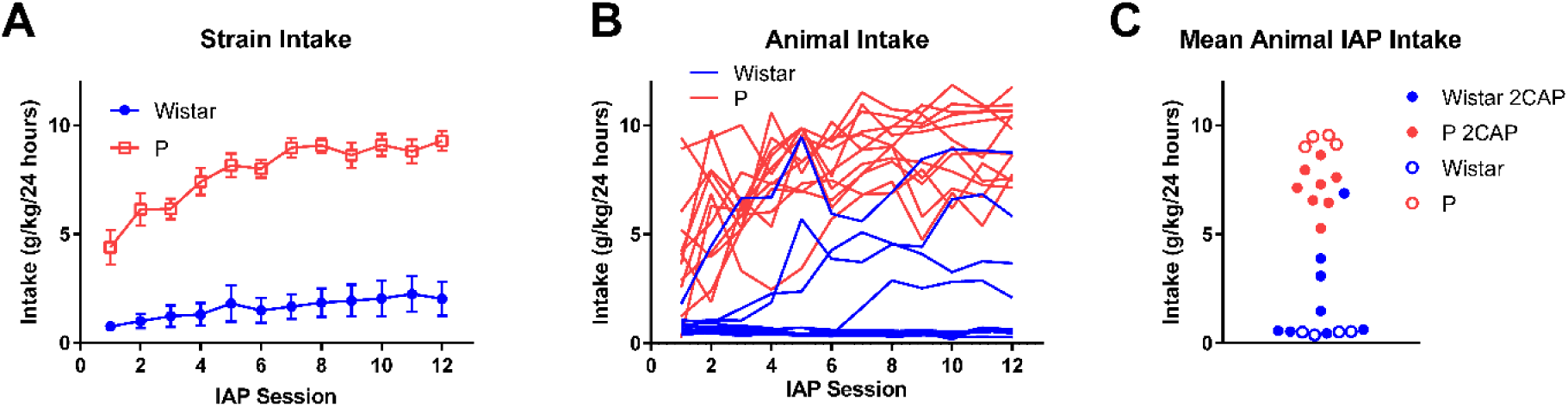
Intermittent access protocol (IAP) consumption. **(A)** As populations, both P rats and Wistars increased consumption throughout IAP (Mean +/-SEM). **(B)** Individual consumption values throughout IAP for all animals. **(C)** 8 P rats and 8 Wistars were selected for 2CAP training (filled circles).

Following IAP, 8 P rats and 8 Wistars were selected for training in the 2CAP task (Figure 2 C). Data for all of these animals is presented throughout the remainder of the analysis. The choice of 8 P rats and 8 Wistars was made based on available training chambers for 2CAP and the desire to maintain balanced numbers of P rats and Wistars. The highest drinking Wistars and lowest drinking P rats throughout IAP were selected to bring strain mean intake closer together. One Wistar with very low intake was selected for training prior to the discovery leaks in the IAP intake data. These leak data points have been manually corrected in these analyses (see analysis code).

### 2CAP Training

A simple, limited access session was used to acclimate animals to the training chamber, to the sippers, and to the reinforcer during the first 3 days of 2CAP training. During this session, animals were given access to 15 licks of 10% ethanol on either side of the operant chamber. They had a maximum time limit of 6 minutes to complete this acclimation session (i.e., reach the lick limit on both sippers). At the end of the acclimation session, a regular 2CAP training session began (see below).

Over the 3 acclimation sessions, time to completion decreased in each group, though P rats completed the task faster (Figure 3 A) (main effect of day: F(2,28) = 35.573, p < 10^−6^, main effect of strain: F(1,14) = 7.174, p = 0.018, interaction: F(2,28) = 2.634, p = 0.089). Also, both P rats and Wistars performed their first lick in progressively shorter times (Figure 3 B) (main effect of day: F(2,28) = 10.716, p < 10^−3^, main effect of strain: F(1,14) = 1.743, p = 0.21, interaction: F(2,28) = 1.67, p = 0.21). Finally, both P rats and Wistars increased their number of licks over the 3 acclimation sessions (Figure 3 C) (main effect of day: F(2,28) = 13.528, p < 10^−4^, main effect of strain: F(1,14) = 3.003, p = 0.11, interaction: F(2,28) = 0.759, p = 0.48). Note that some animals were able to obtain more than 30 total licks because they continued to lick as the sipper was being withdrawn. Importantly, these data provide evidence that all subjects were acclimated to the chamber and were willing to drink at least small amounts of 10% ethanol solution.

**Figure 3:**
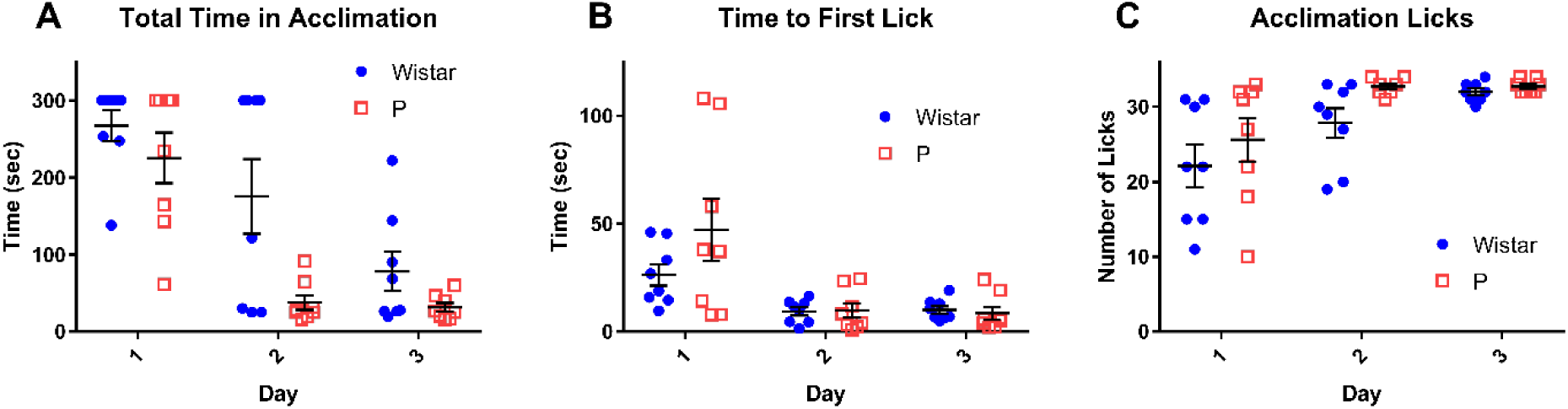
Animals increased licking and lick speed throughout 2CAP acclimation procedure. On the first three days of 2CAP training, animals were acclimated to the test chamber, the sippers, and the reinforcer with up to five minutes of exploration time and 30 total licks, whichever came first. Time required to complete the acclimation task **(A)** and the time to first lick **(B)** decreased over successive acclimation days. **(C)** The number of licks increased throughout acclimation. The sippers withdrew at 30 licks, but animals were frequently able to perform additional licks during sipper withdrawal.

As expected, numerous changes were observed in task performance throughout 2CAP training (Figure 4). During the first 16 days of training, no null trials were included to facilitate task acquisition. Both P rats and Wistars increased alcohol consumption throughout training (Figure 4 A), though P rats reached higher overall consumption levels (main effect of day: F(15,210) = 8.24, p < 10^−6^, main effect of strain: F(1,14) = 27.65, p = 0.0001, interaction: F(15,210) = 1.53, p = 0.096). Both P rats and Wistars increased number of drinking trials throughout training (Figure 4 B), though P rats had more drinking trials (main effect of day: F(15,210) = 5.15, p < 10^−6^, main effect of strain: F(1,14) = 9.17, p = 0.009, interaction: F(15,210) = 0.82, p = 0.65). Both P rats and Wistars decreased the time it took them to drink following sipper insertion (latency to drink) throughout training (Figure 4 C), though P rats exhibited faster and more consistent latencies (one Wistar had no drink trials on the first day which prevented a meaningful latency calculation, main effect of day: F(15,195) = 9.12, p < 10^−6^, main effect of strain: F(1,13) = 9.24, p = 0.009, interaction: F(15,195) = 1.88, p = 0.027). Interestingly, both P rats and Wistars increased the proportion of trials where they approached a sipper following the CS in early stages of training (Figure 4 D), the P rats continued to increase to the point where they approached on nearly all trials (main effect of day: F(15,210) = 7.63, p < 10^−6^, main effect of strain: F(1,14) = 17.46, p = 0.0009, interaction: F(15,210) = 1.30, p = 0.20).

**Figure 4:**
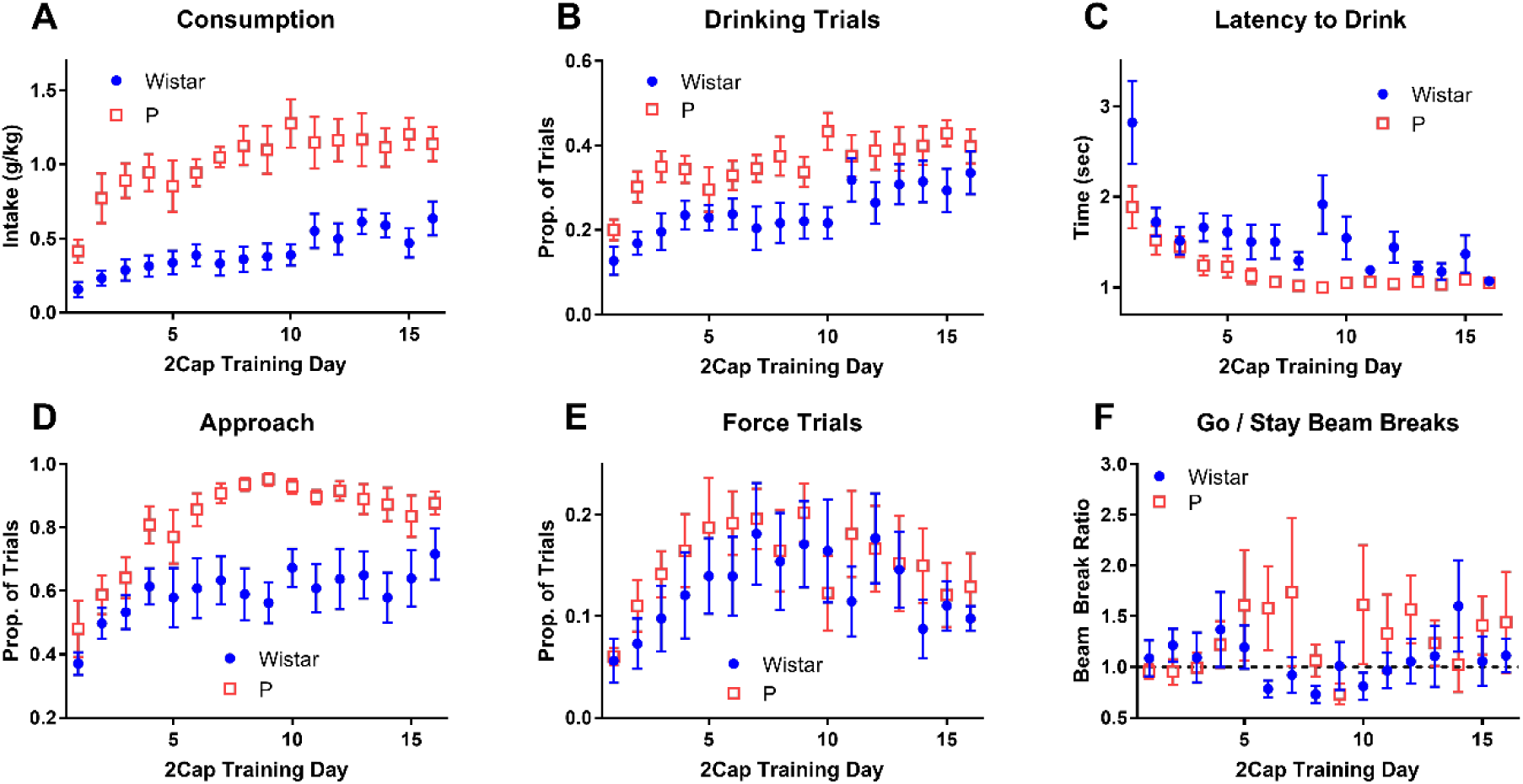
2CAP training prior to null trials. Both P rats and Wistars increased alcohol intake **(A)** and the proportion of trials where the animal drank **(B)**, though P rats did so to larger degrees. **(C)** Latency to first lick following sipper insertion decreased throughout training. **(D)** Both P rats and Wistars increased the proportion of trials where they approached a sipper, but P rats eventually approached on nearly all trials. **(E)** The number of force trials first increased and then decreased throughout training. **(F)** Both P rats and Wistars did not move more on go trials in relation to stay trials.

To ensure that animals would not simple stay near one sipper to obtain access on half the trials, force trials were added such that after three drink trials of one type (stay or go), only trials of the opposite type would appear until the animal drank. The presence of these force trials produced an interesting effect in animal task performance (Figure 4 E). Both P rats and Wistars received increasing numbers of force trials early in training, but then both strains exhibited a decrease in force trials (main effect of day: F(15,210) = 2.12, p = 0.01, main effect of strain: F(1,14) = 1.43, p = 0.25, interaction: F(15,210) = 0.35, p = 0.99). These data indicate that animals initially adopted a strategy wherein they stayed near one sipper and ignored the CS, but that their strategy changed as they learned the CS relationship.

To assess the animals’ use of the CS, the ratio of beam break rates during the go and stay trials was calculated (Figure 4 F). These data were highly variable and exhibited no clear differences between strain or training day (one P rat had a day with no beam breaks on stay trials which prevented a meaningful ratio calculation, main effect of day: F(15,195) = 0.71, p = 0.77, main effect of strain: F(1,13) = 0.78, p = 0.39, interaction: F(15,195) = 1.05, p = 0.40). Furthermore, these data were not consistent above 1, indicating that animals did not move more on go trials relative to stay trial (one sample t-test (comparison mean = 1) for each strain and day produced only 3 out of 32 tests with p < 0.05). These data seem to contradict the force trial data (Figure 4 E) because the animals do not exhibit increased movement for go trials as would expected if they were utilizing the CS to direct their movement.

After 16 days of regular 2CAP training, null trials were added to the task by replacing 10 go and 10 stay trials with 20 null trials (Figure 5). Null trials had identical structure to go and stay trials with the directional tone (4 or 12 kHz) replaced by the attention tone (8 kHz). Both strains’ drink trials tended to be about two-thirds stay trials and one-third go trials prior to the introduction of null trials (Figure 5 A), but the addition of null trials changed this pattern such that all animals drank roughly equally on stay and go trials (main effect of day: F(5,70) = 16.90, p < 10^−6^, main effect of strain: F(1,14) = 0.03, p = 0.86, interaction: F(5,70) = 1.55, p = 0.19). Conversely, the addition of null trials did not change the relationship between movement on stay and go trials for either strain (Figure 5 B) (main effect of day: F(5,70) = 0.33, p = 0.89, main effect of strain: F(1,14) = 0.42, p = 0.53, interaction: F(5,70) = 0.92, p = 0.48). Finally, both P rats and Wistars moved more during null trials than go and stay trials (Figure 5 C) (main effect of day: F(3,42) = 0.16, p = 0.93, main effect of strain: F(1,14) = 0.73, p = 0.41, interaction: F(3,42) = 0.95, p = 0.42, one sample t-test (comparison mean = 0.5) for each strain and day produced 5 out of 8 tests with p < 0.05).

**Figure 5:**
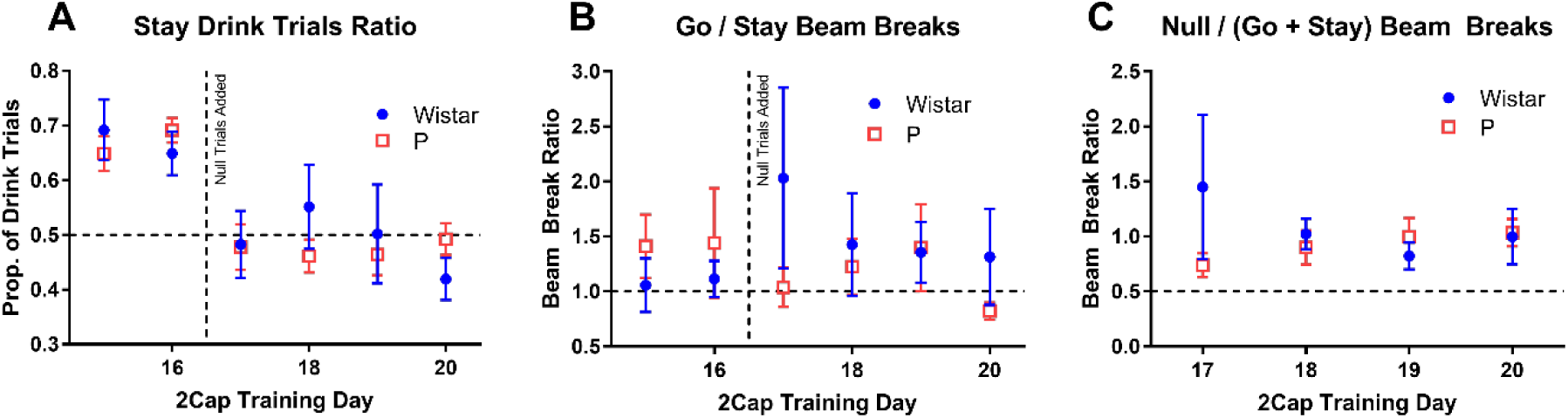
2CAP training with null trials. Starting on the 17^th^ day of 2CAP training, 20 null trials replaced 10 go and 10 stay trials in the 2CAP task. Null trials were identical to go and stay trials, except the directional tone was replaced by continued attention tone frequency and both sippers retracted after being briefly inserted. **(A)** At the end of regular training (days 15 and 16), about 65% of drink trials were stay trials for both P rats and Wistars, but following the introduction of null trials, drinking occurred on roughly balanced trial types for both strains. **(B)** Continuing the pattern from the first 16 days of training, adding null trials did not appear to increase movement to go trials relative to stay trials. **(C)** Both P rats and Wistars appeared to consistently move more on null trials than stay and go trials.

Consumption during IAP and 2CAP were compared to examine if higher consumption animals in IAP also tended to have high consumptions in 2CAP (Figure 6). When data for both strains were fit with a linear regression, a significant positive slope was found, indicating that higher consumption in IAP tended to produce higher consumption in 2CAP (slope +/-SE: 0.072 +/-0.022, R^2^ = 0.44, F(1,14) = 10.8, p = 0.005). However, the large difference in consumption between strains largely drove this effect. When only Wistars (slope +/-SE: 0.021 +/-0.031, R^2^ = 0.07, F(1,6) = 0.43, p = 0.54) or only P rats (slope +/-SE: 0.006 +/-0.081, R^2^ = 0.001, F(1,6) = 0.005, p = 0.95) were examined, neither fit produced a significant slope. Furthermore, when the data were z-scored within each strain and combined (data not shown), the fit did not produce a significant slope (slope +/-SE: 0.144 +/-0.264, R^2^ = 0.021, F(1,14) = 0.296, p = 0.595).

**Figure 6:**
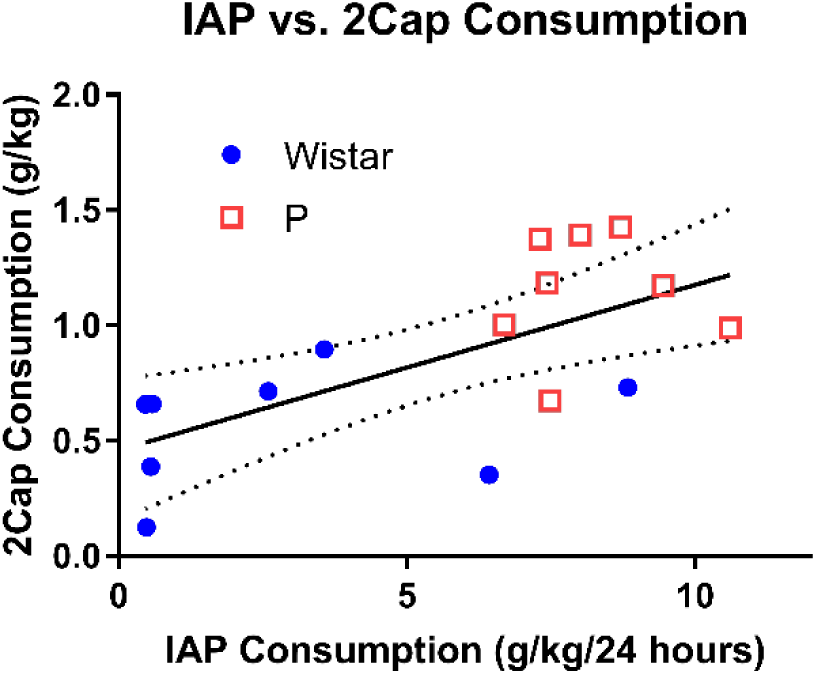
IAP and 2CAP consumption were correlated across all subjects. IAP consumption averaged over last three sessions. 2CAP consumption averaged over last three sessions prior to null trials. Fit shown is for all subjects combined (dashed line: 95% confidence interval).

### Quinine Testing

Aversion resistant drinking was assessed by adulterating the standard 10% ethanol solution with 0.1 g/L quinine. Wistars significantly decreased intake, but P rats did not (main effect of day: F(1,14) = 10.743, p = 0.006; main effect of strain: F(1,14) = 20.9, p = 0.0004; interaction: F(1,14) = 10.15, p = 0.007) (Figure 7 A). However, neither strain significantly reduced the proportion of access trials where the animal drank (i.e., had at least one lick) (main effect of day: F(1,14) = 2.06, p = 0.17; main effect of strain: F(1,14) = 0.72, p = 0.41; interaction: F(1,14) = 1.9, p = 0.19) (Figure 7 B).

**Figure 7:**
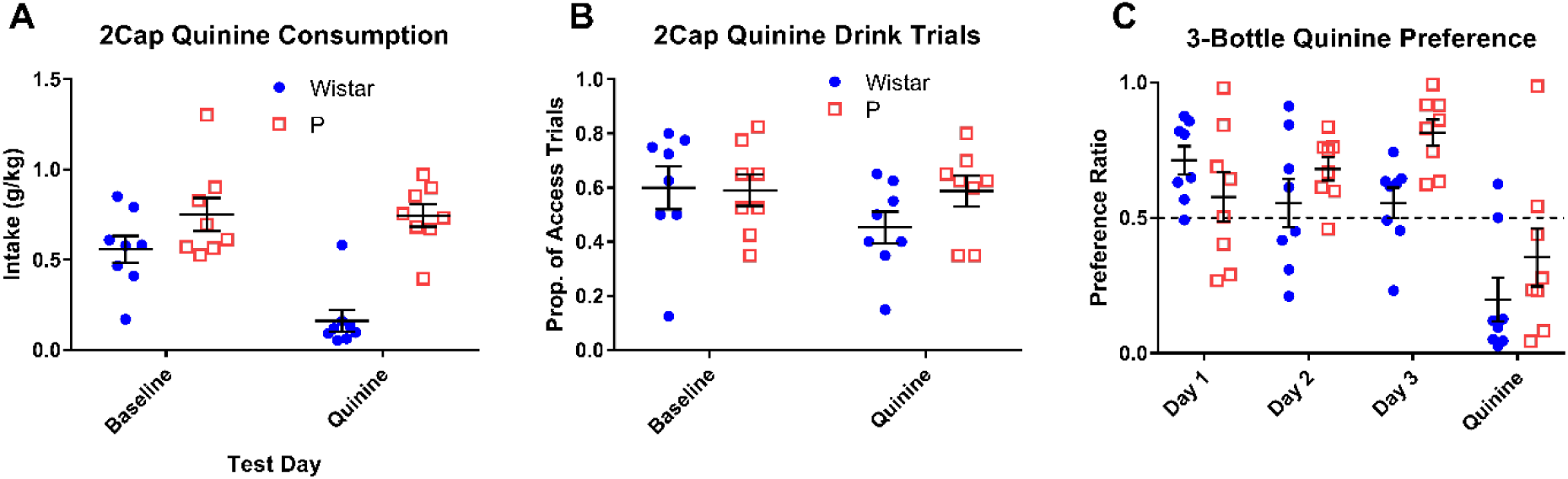
Quinine reduced drinking in Wistars, but not P rats. **(A)** After adulterating the ethanol solution in 2CAP with 0.1 g/L quinine, Wistars showed a significant drop in intake, while P rats showed no change in intake. **(B)** However, we observed only a small change in the number of drink trials (trials where the animal licked at least once) for Wistars, indicated that they continued to test the solution. **(C)** A 3-bottle choice test confirmed that the P rats could taste the quinine following adulteration with 0.1 g/L quinine in the preferred bottle. (Mean +/-SEM in all plots.)

To insure that the P rats could taste the 0.1 g/L quinine concentration in 10% ethanol, a three-bottle (two ethanol bottles, one water bottle) choice, home cage test was conducted (Figure 7 C). After three days of free access to 10% ethanol, the preferred bottle over all three days for each animal was adulterated with 0.1 g/L. The preference ratio (preferred bottle consumption over total consumption) decreased for the quinine bottle for both Wistars and P rats (main effect of day: F(3,14) = 13.01, p < 10^−5^; main effect of strain: F(1,14) = 3.58, p = 0.08; interaction: F(3,14) = 2.59, p = 0.065), indicating that P rats found the 0.1 g/L quinine concentration used in 2CAP testing to be aversive.

### Reversal Testing

To assess the degree to which animals were using the cues to locate alcohol, a tone reversal test session was administered (Figure 8). In this session, the tones for the stay and go cues were reversed for each animal. However, when comparing several performance metrics to behavior during a regular 2CAP session the day before, no significant effects of day were observed. This result indicates that animals were not using the tone to locate alcohol.

**Figure 8:**
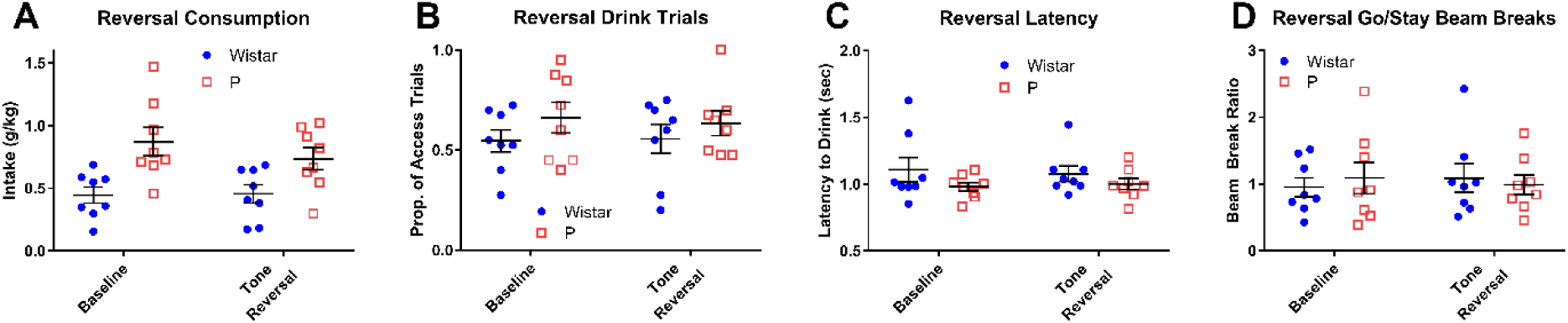
Tone reversal testing indicated animals were not using tone frequencies to locate the correct sipper. For both strains, consumption **(A)**, the proportion of access trials where the animal drank **(B)**, the latency to first lick **(C)**, and the ratio of beam breaks during go vs. stay cues **(D)** did not change when the frequency relationship between go and stay cues was reversed. (Mean +/-SEM in all plots.)

### Free Access

During a free access session, animals were given uninterrupted access to 10% ethanol to assess motivation to consume 10% ethanol solution in the 2CAP setting (Figure 9). During the free access session, the sippers entered the chamber at the beginning and did not retract throughout the entire session (duration matched to regular 2CAP). In comparison to a regular 2CAP session on the previous day, animals consumed more alcohol during the free access session (main effect of day: F(1,14) = 55.45, p < 10^−5^). In addition, P rats consumed more than Wistars (main effect of strain: F(1,14) = 11.69, p = 0.004) and an interaction between strain and day was observed (F(1,14) = 5.44, p = 0.035). This large increase in consumption in the free access session indicates that both P rats and Wistars are motivated to consume alcohol during the regular 2CAP task.

**Figure 9:**
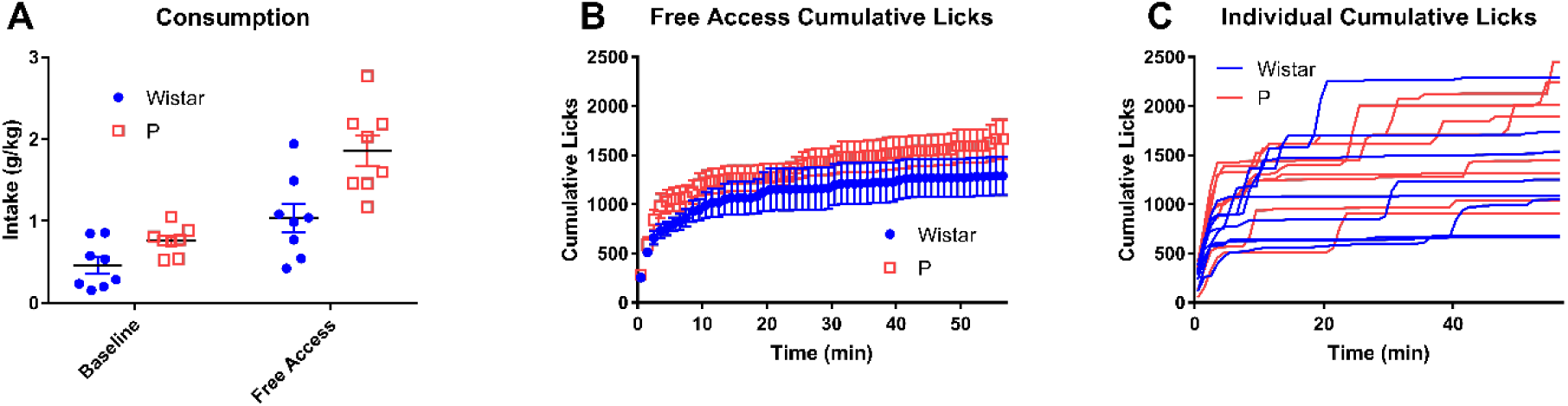
Free access testing indicated that animals were motivated to drink during standard 2CAP. Free Access testing was performed by giving the animals free access to ethanol in their standard 2CAP test chambers for the same overall period of time as a standard 2CAP session. **(A)** Intake increased from standard 2CAP (baseline, one day earlier) to Free Access test. **(B)** Cumulative lick distributions for strains (mean +/-SEM) and **(C)** individual subjects showed that animals performed most of their consumption during the first few minutes of the session with subsequent drinking occurring in fairly discrete bouts.

### BEC Data

Blood ethanol concentration (BEC) measurements were taken immediately following a regular 2CAP session (Figure 10 A) and a free access 2CAP session (Figure 10 B). A fit of the regular 2CAP intake vs. BEC for all animals found a significant slope (mean +/-SE: 80.7 +/-33.2, F(1,14) = 5.92, p = 0.029) (Figure 10 A). Similarly, a fit of the free access intake vs. BEC for all animals also found a significant slope (mean +/-SE: 44.7 +/-8.1, F(1,14) = 30.3, p < 10^−4^) (Figure 10 B). The relationships between intake and BEC for these two types of tasks were different due to differences in drinking patterns. The regular 2CAP required animals to spread drinking out throughout the task due to regular intervals of access during trials, whereas the free access session produced drinking patterns with large bouts clustered near the beginning of the session (Figure 9 B).

**Figure 10:**
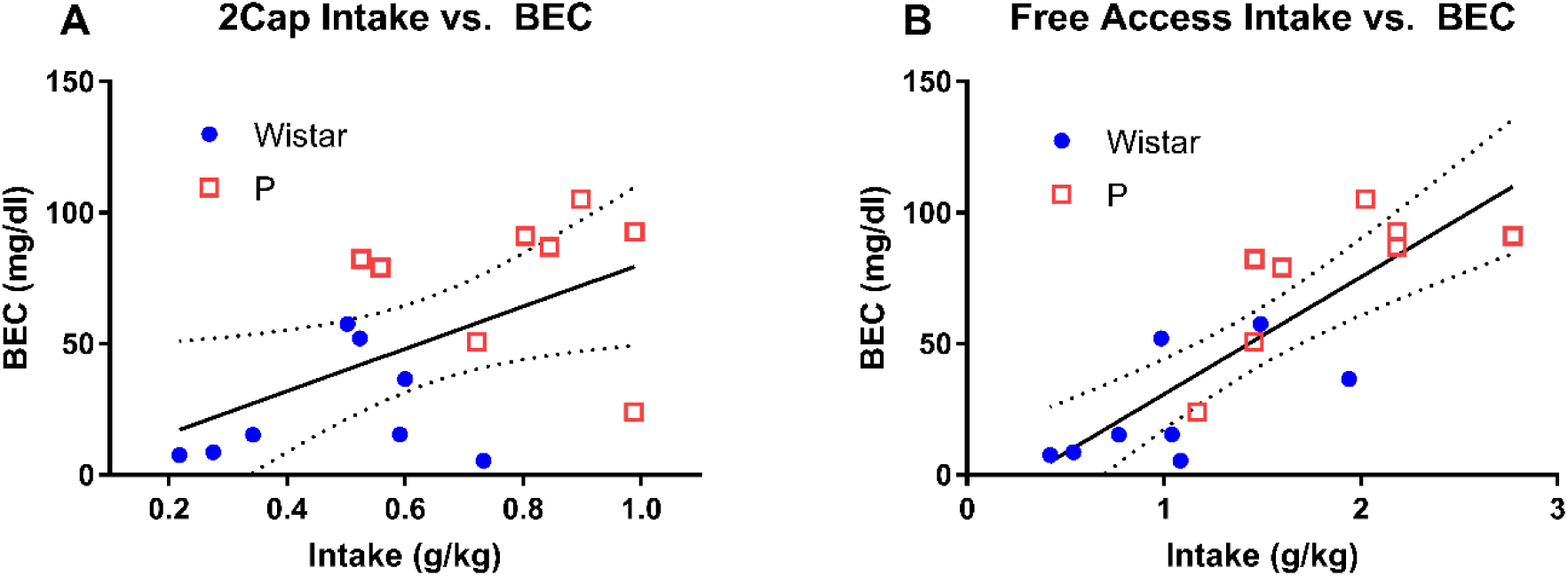
BEC results indicated that animals experienced the pharmacological effects of ethanol in both standard 2CAP and Free Access testing. **(A)** Immediately following 2CAP, 2 Wistars (total: 8) and 7 P rats (total: 8) achieved BEC values greater than 40 mg/dl. **(B)** Immediately following Free Access testing, the same number of animals achieved BEC values greater than 40 mg/dl. During Free Access testing most animals consumed the majority of the alcohol during the first few minutes of the session (i.e., roughly one hour prior to blood draw), whereas during standard 2CAP animals were forced to more evenly disperse consumption throughout the session. These differences in consumption pattern resulted in different intake vs. BEC relationships.

## Discussion

### Aversion Resistant Drinking

We used quinine adulteration to assess aversion resistant drinking (Hopf et al., 2010) in alcohol preferring P rats and Wistars (see Quinine Testing). We found that P rats did not reduce intake when drinking quinine adulterated alcohol in the 2CAP task, but Wistars did reduce intake. Importantly, when given the option to drink quinine adulterated alcohol or non-adulterated alcohol in free access home cage drinking, these P rats preferred non-adulterated alcohol, indicating that they found this concentration of alcohol aversive. Also, though Wistars reduced intake in the 2CAP task with quinine adulterated alcohol, they did not reduce number of drink trials, indicating that they were motivated to consume alcohol in the 2CAP task. Overall, these results indicate that drinking by P rats in this task was inflexible and aversion resistant, whereas Wistars maintained control over drinking and were able to modify their drinking pattern based on the aversive stimuli.

These results are important because they demonstrate that these two strains of rats can serve as models of aversion resistant, inflexible drinking and aversion sensitive, flexible drinking. In the future, we will investigate the cause of these differences between P rats and Wistars. Though we showed that P rats found this concentration of quinine aversive, perhaps it is only less aversive to P rats than Wistars. Indeed, several dose, strain, and species dependent effects have been observed with quinine adulteration as a model of aversion resistant drinking. For instance, other authors have shown that with enough drinking history and a lower dose of quinine, Wistar rats will exhibit aversion resistant drinking (Seif et al., 2013). Also, it has been shown that a single alcohol session with C57/BL6 mice produced aversion resistant drinking (Lei, Wegner, Yu, Simms, & Hopf, 2016). We would expect that there is some higher dose of quinine that would render P rats quinine sensitive, but we have not tested for such a dose. However, understanding why P rats are aversion resistant and Wistars are aversion sensitive is an important question because it mirrors the vital question of why some people continue to drink despite negative consequences, while others do not.

The neurological differences underlying aversion-resistant drinking are only beginning to be understood. In a particularly important study, it was show that connections from medial prefrontal cortex to ventral striatum are necessary for aversion-resistant drinking (Seif et al., 2013). Numerous molecular changes have been observed in amygdala following chronic-intermittent alcohol exposure (Hopf & Lesscher, 2014), including gene expression factors that regulate ARD (Lesscher, Houthuijzen, Groot Koerkamp, Holstege, & Vanderschuren, 2012). Finally, disruption of neural activity in insular cortex (insula) has been shown to reduce ARD (Chen & Lasek, 2018). Finally, recent behavioral evidence for a more automated “head down and push” strategy in aversion resistant consumption (Darevsky et al., 2018) indicates subtle behavioral differences in consumption that may be mediated by different neural circuits. In the future, we hope to investigate how aversion resistant and aversion sensitive drinking arise neurologically in P rats and Wistars in order to elucidate possible causes of aversion resistant drinking in humans.

### Free Access Drinking

By examining the distribution lick times during free access drinking, we observed several important features of the animals’ drinking patterns (see Free Access). First, both P rats and Wistars increased drinking in free access relative to regular 2CAP drinking. This indicates that the both strains were motivated to consume alcohol and were limited to less than their free access consumption levels in the 2CAP task. Second, drinking in free access occurred primarily in the first few minutes of the session and in discrete bouts. Therefore, the animals tended to quickly drink to a certain threshold immediately following the beginning of alcohol access and then tended to drink only sporadically throughout the remainder of the session. As such, this task could provide a useful model to assess the neurobiological and behavioral processes that underlie front-loading and maintenance drinking.

### 2CAP Task

The audio 2CAP task used in this study was adopted from a previously published visual version of the 2CAP task (Linsenbardt & Lapish, 2015; Linsenbardt et al., 2018; McCane et al., 2014). However, in the current task, evidence for an association with the CS+ and a specific command was not detected (go vs. stay) (Figure 4 F) as animals did not change their behavior during the null CS (Figure 5 C).

There are at least two possible explanations for these results. First, it is possible that with further training, the animals would eventually learn the CS direction and to ignore the null CS. Previous research has shown that Wistar rats can differentiate between similar frequencies to those used in this task (Ono, Kudoh, & Shibuki, 2006), but perhaps these tones are not salient enough to generate a change in behavior in the training period tested. Second, it is possible that, given the large amount of access available to each animal without learning the CS direction association, the animals were not motivated to learn this association. Furthermore, because there was no cost to exploring for alcohol following the null CS, it is possible that the animals will never fully extinguish searching behavior during the null CS.

### Genetic Risk

The data presented herein indicate that following about 8 weeks of alcohol exposure, P rats are resistant to quinine devaluation of alcohol drinking whereas Wistar rats are not. These data may indicate that genetic risk for excessive drinking accelerates the acquisition of quinine resistance. However, asymmetries in alcohol consumption history between P rats and Wistars complicate this interpretation. Future work will be required to clearly parse the influence of genetic risk and alcohol consumption history in quinine resistance.

## References

1. Beckwith, S. W., & Czachowski, C. L. (2014). Increased delay discounting tracks with a high ethanol-seeking phenotype and subsequent ethanol seeking but not consumption. Alcoholism, Clinical and Experimental Research, 38(10), 2607–2614. doi:10.1111/acer.12523

2. Beckwith, S. W., & Czachowski, C. L. (2016). Alcohol-preferring P rats exhibit elevated motor impulsivity concomitant with operant responding and self-administration of alcohol. Alcoholism, Clinical and Experimental Research, 40(5), 1100–1110. doi:10.1111/acer.13044

3. Bell, R. L., Rodd, Z. A., Lumeng, L., Murphy, J. M., & McBride, W. J. (2006). The alcohol-preferring P rat and animal models of excessive alcohol drinking. Addiction Biology, 11(3-4), 270–288. doi:10.1111/j.1369-1600.2005.00029.x

4. Chen, H., & Lasek, A. W. (2018). Perineuronal nets in the insula regulate aversion-resistant alcohol drinking. bioRxiv, 504571. doi:10.1101/504571

5. Darevsky, D., Gill, T. M., Vitale, K. R., Hu, B., Wegner, S. A., & Hopf, F. W. (2018). Drinking despite adversity: behavioral evidence for a head down and push strategy of conflict-resistant alcohol drinking in rats. Addiction Biology, 24, 426–437. doi:10.1111/adb.12608

6. Edenberg, H. J., & Foroud, T. (2013). Genetics and alcoholism. Nature Reviews Gatroenterology and Hepatology, 10(8), 487–494. doi:10.1038/nrgastro.2013.86

7. Engleman, E. A., Ingraham, C. M., McBride, W. J., Lumeng, L., & Murphy, J. M. (2006). Extracellular dopamine levels are lower in the medial prefrontal cortex of alcohol-preferring rats compared to Wistar rats. Alcohol, 38(1), 5–12. doi:10.1016/j.alcohol.2006.03.001

8. Enoch, M. A. (2013). Genetic influences on the development of alcoholism. Current Psychiatry Reports, 15(11). doi:10.1007/s11920-013-0412-1

9. Field, M., & Cox, W. M. (2008). Attentional bias in addictive behaviors: a review of its development, causes, and consequences. Drug and Alcohol Dependence, 97(1-2), 1–20. doi:10.1016/j.drugalcdep.2008.03.030

10. Gilpin, N. W., Stewart, R. B., & Badia-Elder, N. E. (2008). Neuropeptide Y administration into the amygdala suppresses ethanol drinking in alcohol-preferring (P) rats following multiple deprivations. Pharmacology, Biochemistry and Behavior, 90(3), 470–474. doi:10.1016/j.pbb.2008.04.005

11. Hopf, F. W., Chang, S. J., Sparta, D. R., Bowers, M. S., & Bonci, A. (2010). Motivation for alcohol becomes resistant to quinine adulteration after 3 to 4 months of intermittent alcohol self-administration. Alcoholism, Clinical and Experimental Research, 34(9), 1565–1573. doi:10.1111/j.1530-0277.2010.01241.x

12. Hopf, F. W., & Lesscher, H. M. B. (2014). Rodent models for compulsive alcohol intake. Alcohol, 48(3), 253–264. doi:10.1016/j.alcohol.2014.03.001

13. Kreusch, F., Vilenne, A., & Quertemont, E. (2013). Response inhibition toward alcohol-related cues using an alcohol go/no-go task in problem and non-problem drinkers. Addictive Behaviors, 38(10), 2520–2528. doi:10.1016/j.addbeh.2013.04.007

14. Lei, K., Wegner, S. A., Yu, J. H., Simms, J. A., & Hopf, F. W. (2016). A single alcohol drinking session is sufficient to enable subsequent aversion-resistant consumption in mice. Alcohol, 55, 9–16. doi:10.1016/j.alcohol.2016.07.008

15. Lesscher, H. M. B., Houthuijzen, J. M., Groot Koerkamp, M. J., Holstege, F. C. P., & Vanderschuren, L. J. M. J. (2012). Amygdala 14-3-3ζ. PloS One, 7(5), e37999. doi:10.1371/journal.pone.0037999

16. Linsenbardt, D. N., & Lapish, C. C. (2015). Neural firing in the prefrontal cortex during alcohol intake in alcohol preferring ‘P’ vs. Wistar rats. Alcoholism, Clinical and Experimental Research, 39(9), 1642–1653. doi:10.1111/acer.12804

17. Linsenbardt, D. N., Smoker, M. P., Janetsian-Fritz, S. S., & Lapish, C. C. (2016). Impulsivity in rodents with a genetic predisposition for excessive alcohol consumption is associated with a lack of a prospective strategy. Cognitive, Affective, & Behavioral Neuroscience, 17(2), 235–251. doi:10.3758/s13415-016-0475-7

18. Linsenbardt, D. N., Timme, N. M., & Lapish, C. C. (2018). Encoding of the intent to drink alcohol by the prefrontal cortex is blunted in rats with a family history of excessive drinking. bioRxiv. doi:10.1101/490664

19. McBride, W. J., Rodd, Z. A., Bell, R. L., Lumeng, L., & Li, T. K. (2014). The alcohol-preferring (P) and high-alcohol-drinking (HAD) rats - animal models of alcoholism. Alcohol, 48, 209–215. doi:10.1016/j.alcohol.2013.09.044

20. McCane, A. M., Czachowski, C. L., & Lapish, C. C. (2014). Tolcapone suppresses ethanol intake in alcohol preferring rats performing a novel cued access protocol. Alcoholism, Clinical and Experimental Research, 38(9), 2468–2478. doi:10.1111/acer.12515

21. Ono, K., Kudoh, M., & Shibuki, K. (2006). Roles of the auditory cortex in discrimination learning by rats. European Journal of Neuroscience, 33(1-2), 1623–1632. doi:10.1016/S0378-5955(98)00162-2

22. Sanchis-Segura, C., & Spanagel, R. (2006). Behavioural assessment of drug reinforcement and addictive features in rodents: an overview. Addiction Biology, 11(1), 2–38. doi:10.1111/j.1355-6215.2006.00012.x

23. Seif, T., Chang, S. J., Simms, J. A., Gibb, S. L., Dadgar, J., Chen, B. T., … Hopf, F. W. (2013). Cortical activation of accumbens hyperpolarization-active NMDARs mediates aversion-resistant alcohol intake. Nature Neuroscience, 16(8), 1094–1100. doi:10.1038/nn.3445

24. Simms, J. A., Steensland, P., Medina, B., Abernathy, K. E., Chandler, L. J., Wise, R., & Bartlett, S. E. (2008). Intermittent access to 20% ethanol induces high ethanol consumption in Long-Evans and Wistar rats. Alcoholism: Clinical and Experimental Research, 32(10), 1816–1823. doi:10.1111/j.1530-0277.2008.00753.x

25. Wiers, C. E., Stelzel, C., Park, S. Q., Gawron, C. K., Ludwig, V. U., Gutwinski, S., … Bermpohl, F. (2014). Neural correlates of alcohol-approach bias in alcohol addiction: the spirit is willing but the flesh is weak for spirits. Neuropsychopharmacology, 39, 688–697. doi:10.1038/npp.2013.252

26. Zhou, Z., Karlsson, C., Liang, T., Xiong, W., Kimura, M., Tapocik, J. D., … Goldman, D. (2013). Loss of metabotropic glutamate receptor 2 escalates alcohol consumption. PNAS, 110(42), 16963–16968. doi:10.1073/pnas.1309839110

